# Solid-state nanopore sensing reveals conformational changes induced by a mutation in a neuron-specific tRNA^Arg^

**DOI:** 10.1101/2025.04.08.647894

**Authors:** Shankar Dutt, Lien B. Lai, Rahul Mehta, Buddini I. Karawdeniya, Y.M. Nuwan D.Y. Bandara, Andrew J. Clulow, Sebastian Glatt, Venkat Gopalan, Patrick Kluth

**Affiliations:** Department of Materials Physics, Research School of Physics, Australian National University, Canberra, ACT 2601, Australia; Department of Chemistry and Biochemistry, Center for RNA Biology, The Ohio State University, Columbus, Ohio 43210, USA; Malopolska Centre of Biotechnology, Jagiellonian University, 30-387, Krakow, Poland; Doctoral School of Exact and Natural Sciences, Jagiellonian University, 30-348, Krakow, Poland; Department of Electronics Materials Engineering, Research School of Physics, Australian National University, Canberra, ACT 2601, Australia; Research School of Chemistry, Australian National University, Canberra, ACT 2601, Australia; Australian Synchrotron, ANSTO, 800 Blackburn Road, Clayton, VIC 3168, Australia; University of Veterinary Medicine Vienna, 1210 Vienna, Austria; Department of Biomedical Engineering, The Ohio State University, Columbus, Ohio 43210, USA; Department of Chemistry and Biochemistry, The Ohio State University, Columbus, Ohio 43210, USA

**Keywords:** Solid-state nanopore sensing, single-molecule analysis, tRNA structure, conformational sampling, structural plasticity, SAXS, cryo-EM, time-resolved measurements

## Abstract

We demonstrate that solid-state nanopore sensing is a powerful single-molecule method for analyzing RNA conformational ensembles. As a model, we employed n-Tr20, a neuron-specific cytoplasmic tRNA^Arg^_UCU_, whose C50U mutation is associated with neurodegeneration in C57BL/6J mice. Maturation of the Tr20^C50U^ precursor is impaired as the mutation stabilizes a conformational ensemble different from the wild-type. To gain insights into how this mutation engenders structural differences, we used solid-state nanopore sensing for the real-time identification of metastable conformers that are not easily observable by ensemble methods. Ion-current traces recorded using an 8-nm nanopore revealed broad contours of the conformational landscape of n-Tr20/n-Tr20^C50U^ ± Mg^2+^. Additionally, cryo-EM analysis and small-angle X-ray scattering studies revealed structural plasticity even more than predicted from the nanopore-sensing data. Since dynamics undergird RNA (dys)function in cellular physiology and pathology, nanopore sensing to determine RNA conformational sampling is a valuable addition to the growing RNA structural analysis toolkit.

**Graphical Abstract:** 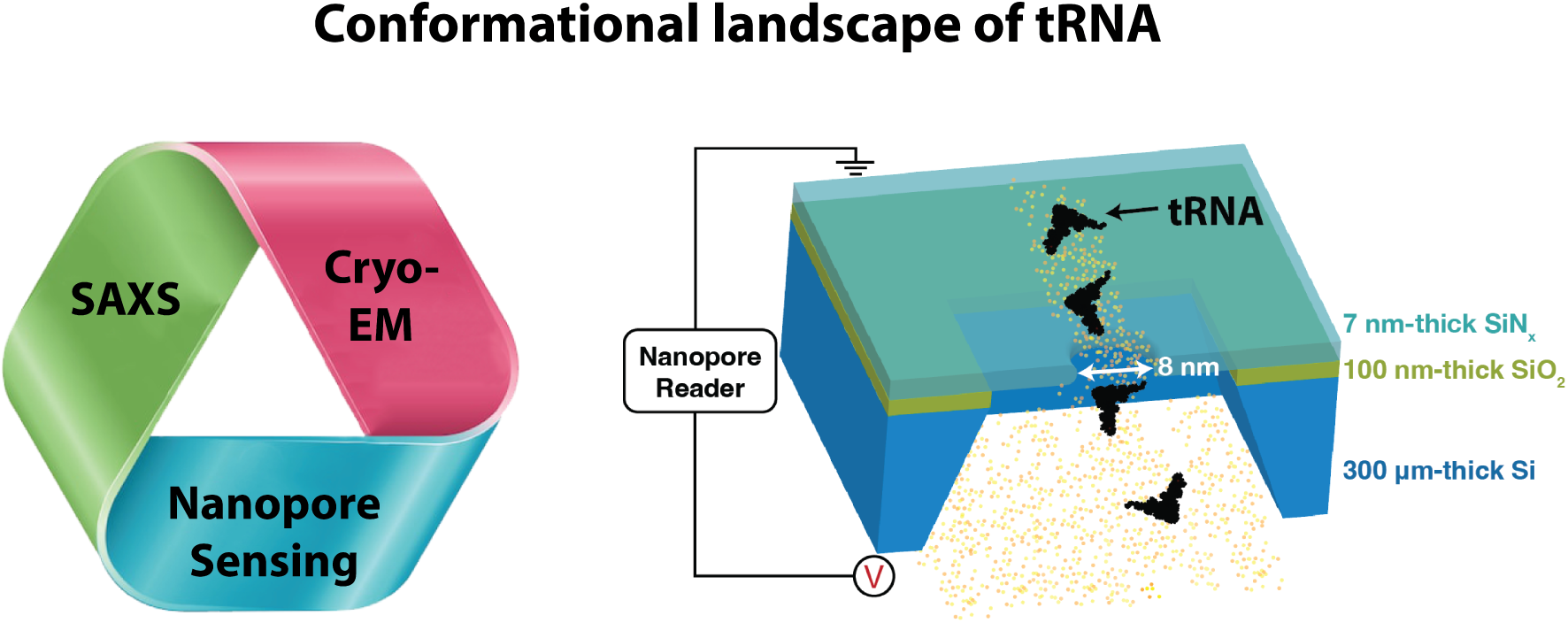

## Introduction

The folding trajectory of structured RNAs typically entails a rugged energy landscape defined by several local minima, which reflect potential kinetic traps en route to a native fold [**1**]. Although chaperones and metal ions lower the energy barriers between intermediates and help convert metastable states to the final native structure, real-time observation of this conformational sampling has been largely intractable using conventional analytical methods. To address this limitation, we explore a nanopore-sensing approach and showcase its value with a transfer RNA (tRNA) as the model.

The most abundant cellular RNA (in molar amounts) is tRNA. These small non-coding RNAs (∼75 nt) deliver amino acids attached to their 3*^′^* end to the ribosomes for protein synthesis [**1**]. Although often overlooked as a cause of diseases due to gene redundancy, there is increased appreciation that dysregulation of individual tRNAs could lead to neurodegenerative diseases, developmental defects, cancer, and hearing loss [**2**–**6**]. Given the extensive suite of post-transcriptional modifications (total ∼100, average of 13 per mammalian tRNA), this fine-tuning of tRNA structure and function in response to changing levels of cellular metabolites provides another tier of regulation [**7**–**11**]. However, understanding how genetic mutations or modifications influence RNA structural dynamics remains a formidable challenge, given the intrinsic propensity of RNAs to sample stable inter/intramolecular interactions and adopt alternative structures [**12**–**14**].

Traditionally, tRNAs have been studied employing ensemble methods, such as nuclear magnetic resonance spectroscopy [**15**, **16**] and X-ray crystallography [**17**]. Although these techniques have provided high-resolution structures, they do not capture metastable states essential to understanding the tRNA structural dynamics, especially under different physiological conditions. On the other hand, single-molecule techniques such as single-molecule fluorescence resonance energy transfer (smFRET) and optical tweezer-based methodologies show promise for characterizing RNA dynamics, but their low-throughput precludes studies of large molecular populations [**18**].

Solid-state nanopore-enabled resistive pulse sensing (**Fig. 1a**) offers label-free monitoring of single-molecule biomolecular dynamics providing time-resolved structural information of the biomolecules and, as shown in the present study, could potentially become a valuable technique for studying tRNA conformational changes. By bridging the gap between detailed single-molecule insights and high-throughput data from hundreds of thousands of molecules (up to 10,000 translocation events/min [**19**]), solid-state nanopore sensing offers the advantages of both approaches, enabling robust statistical analysis and detailed conformational understanding at the level of individual molecules.

**Figure 1:**
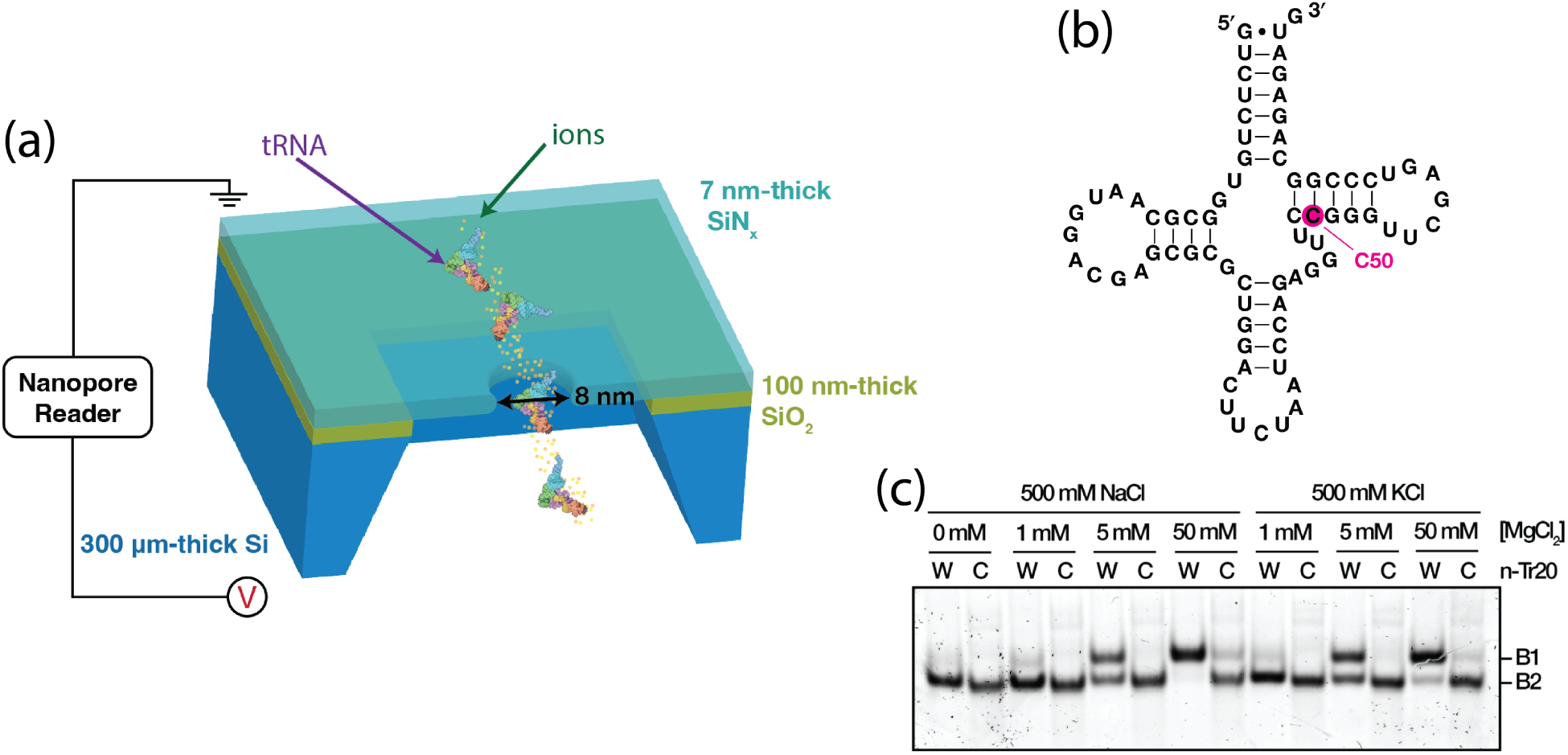
Use of a solid-state nanopore sensor and native PAGE to study tRNA conformational dynamics. (a) Schematic of tRNA translocation through a 7-nm-thick SiN*_x_* membrane with a 8-nm diameter nanopore. A 100-nm-thick SiO_2_ underlayer was used to reduce dielectric noise [**19**, **53**]. The layers are supported by a 300-*µ*m thick silicon wafer. (b) Secondary structure of n-Tr20 with the C50U mutation depicted. (c) Native PAGE analysis of n-Tr20 (W) and Tr20^C50U^ (C) folded in the indicated concentrations of MgCl_2_ and either 500 mM NaCl or KCl. The tRNAs separated into two main bands, denoted as B1 and B2 (see text for details). It is likely that these bands represent conformational ensembles and not distinct species.

A solid-state nanopore is a nanofluidic device comprising a nanometer-scale channel that connects two electrolyte reservoirs. These nanochannels have been fabricated in insulating materials such as silicon nitride (SiN_x_) [**19**–**21**], silicon dioxide (SiO_2_) [**22**, **23**], hafnium oxide (HfO_x_) [**24**, **25**], polymers [**26**, **27**] or two-dimensional (2D) materials such as graphene[**28**, **29**], and molybdenum disulfide [**30**]. Under an applied electro-potential bias, the flow of ions through the nanopore generates a baseline current. An event occurs when a (bio)molecule translocates through the pore and perturbs the ion flow, thereby causing current fluctuations that in turn are recorded as resistive or conductive pulses characteristic of the (bio)molecule. This current signature encodes information about the structure of the translocating (bio)molecule. Recent advances in thin-membrane fabrication by controlled etching or thinning [**19**, **31**, **32**], nanopore fabrication by the controlled breakdown method [**33**–**36**], and the use of machine learning [**37**–**41**] have paved the way to widespread adoption of solid-state nanopores for sensing and structural analysis of different molecules.

Solid-state nanopores (**Fig. 1a**) have been successfully used to probe the conformations of proteins and their complexes [**37**, **42**, **43**] and study the flexibility/rigidity of enzymes [**44**]. Recently, solid-state nanopores have also shown promise in sensing RNA molecules [**45**, **46**]. These studies have yielded information on structure-function correlations that extend beyond those obtained from static crystallographic snapshots. Here, we employed nanopore sensing to study the structural changes of n-Tr20, a neuron-specific tRNA^Arg^ whose C50U mutation is associated with neurodegeneration in C57BL/6J mice [**5**].

We investigated Mg^2+^-dependent and time-resolved conformational dynamics of n-Tr20 and n-Tr20^C50U^. To gain additional insights into the wide conformational space sampled by n-Tr20 and n-Tr20^C50U^, we performed synchrotron-based small-angle X-ray scattering (SAXS) and employed cryo-EM analysis to complement our solid-state nanopore measurements. By capturing data from ∼ 3 × 10^6^ tRNAs, we show how solid-state nanopore-based resistive pulse sensing and analysis surpasses the throughput limitation of existing approaches and is a valuable tool to map the temporal structural properties of tRNAs and potentially other RNAs and RNA-protein complexes.

## Materials & Methods

### Preparation of mature n-Tr20 and n-Tr20^C50U^

The mature tRNA sequences were cloned downstream of the T7 RNA polymerase promoter in pBT7 as described in our earlier work [**14**]. To generate transcripts by run-off in vitro transcription using T7 RNA polymerase, the DNA templates were amplified by PCR using the above clones and the primers F-ext (5*^′^*-CGACGTTGTAAAACGACGGCCAG-3*^′^*) and pARGnoCCA-R (5*^′^*-CATCTCT GCCGGGACTCGAAC-3*^′^*). The transcripts were extensively dialyzed against water and then precipitated with sodium acetate and ethanol. After resuspension in water, the final RNA concentration was determined using a NanoDrop spectrophotometer and the specific extinction coefficient of the RNA at 260 nm.

### Nanopore fabrication

Silicon nitride windows of 40 *µ*m × 40 *µ*m in size and of ∼ 7 nm thickness with a silicon dioxide underlayer of 100 nm thickness were fabricated using methods described before [**19**]. Nanopores in these windows were fabricated using the controlled breakdown method [**19**, **34**] by applying *<*1 V/nm voltage across the membrane. One molar KCl (pH 7) was used for nanopore fabrication. The voltage application ceased immediately upon the detection of a sharp increase in current, signaling the successful creation of a nanopore. To determine the dimensions of the crafted nanopore, we analyzed its open pore conductance. This objective was achieved by performing a linear fit of the current-voltage data, which were recorded using the eNPR-200 amplifier. The diameter of the pore was calculated using the following equation [**19**, **47**]:

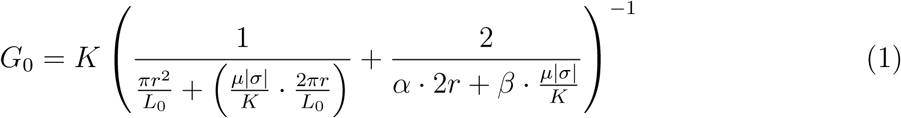

where *G*_0_, *r*, *K*, *σ*, *µ*, *α*, and *β* are, respectively, the open-pore conductance, nanopore radius, electrolyte conductivity, nanopore surface charge density, mobility of counterions proximal to the surface, and model-dependent parameters (both *α* and *β* are set to 2).

### Nanopore data acquisition and analysis

The translocation measurements of the tRNA were performed in 5 mM Tris-HCl (pH 8) and 500 mM NaCl under an applied bias of 400 mV. Before translocation experiments, tRNA was refolded in water (95 °C for 3 min, 37 °C for 10 min) prior to addition of the electrolyte at the specified concentration. The samples were incubated for 30 min and the measurements were performed at ∼22 °C. Experiments were carried out after the incubation period and the time was set as t_0_ when MgCl_2_ was added (in the case of time-resolved experiments). 10 nM tRNA was used for translocation experiments. Data was acquired at 200 kHz sampling rate and low-pass filtered using a 35 kHz Butterworth filter [**37**]. For event extraction, we used the methodology detailed in our previous work, applying threshold values of at least five times the standard deviation of the baseline current in the analysis window [**37**]. For generating time-resolved graphs, the data were segmented into 7-minute intervals, since this duration typically yielded approximately 10,000 events. For the analysis of these events, the histogram plots were subjected to Gaussian fitting based on machine learning to deconvolute data.

### Native polyacrylamide gel electrophoresis (nPAGE)

10 pM of either n-Tr20 or n-Tr20^C50U^ in 8 *µ*L water was incubated at 95 °C for 3 min and at 37 °C for 10 min before adding 2 *µ*L of 5x buffer [5x Nanopore Analysis (Fig. 1c): 25 mM Tris-HCl (pH 8 at 22°C), 2.5 M NaCl or KCl; 5x Folding and 5x Cryo-EM (Fig. **S6**): 25 mM HEPES (pH 7.4 at 22°C) plus 100 mM NaCl or 250 mM each of NaCl and KCl, respectively; each buffer also contained varying amounts of MgCl_2_ as indicated.]. The sample was then incubated at 37 °C for an additional 30 min. Subsequently, 3 *µ*L of loading buffer [50% (v/v) glycerol, 0.05% (w/v) bromophenol blue, 0.05% (w/v) xylene cyanol] was added. TBM buffer (89 mM Tris base, 89 mM boric acid, 2 mM MgCl_2_) was used to prepare a 5% (w/v) polyacrylamide (19:1 acrylamide:bis-acrylamide) gel and used as the running buffer for electrophoresis. The gels (17 cm width × 15 cm height) were electrophoresed at 180 V and 4 °C until the bromophenol blue dye reached the bottom. Gels were then stained with SYBR Gold (Invitrogen) in TBE buffer for 5 min and then scanned using a Typhoon phosphorimager (GE Healthcare) and the Cy2 imaging mode.

### Small-angle X-ray scattering (SAXS)

Consistent with the conditions of our nanopore experiments, 0.5 mg/mL of mature tRNA in water was incubated at 95°C for 3 minutes, followed by cooling at 37 °C for 5 minutes. Subsequently, 5 mM Tris-HCL (pH 8) and 500 mM NaCl were added, along with 0 mM or 50 mM MgCl_2_. Transmission SAXS measurements were performed on the bioSAXS beamline of the Australian Synchrotron using X-rays with an energy of 12.4 keV, corresponding to a wavelength of 1 Å. Data collection was performed with an automated sample changer (CoFlow) and a Pilatus3X-2M detector (in a vacuum-movable detector system) with a pixel size of 172 *µ*m^2^. The sample-to-detector distance was calibrated at 3156 mm against a combination of silver behenate and LUDOX^®^ colloidal silica particles used as a secondary standard. To mitigate radiation damage, a laminar sheath flow of the buffer solution was used along the capillary walls. Each sample (50 *µ*L) underwent 20 acquisitions, each lasting 0.5 seconds throughout the injection of the sample. Data reduction was performed using the pyFAI library[**48**] with a custom-written script. The reduced data were then averaged and buffer-only measurements subtracted. The resulting scattering curves were analyzed using the ATSAS software suite[**49**]. To determine the radius of gyration, the linear region of *ln*(I) *vs* q^2^ was identified (qR*_g_ <* 1.3) for each curve using Guinier analysis.

### Electron microscopy

QUANTIFOIL^®^ R 2/1 copper grids (200 mesh) were glow discharged using a Leica EM ACE 200 glow discharger (8 mA, 60 s). 3 *µ*L of 34 *µ*M (n-Tr20) or 54 *µ*M (n-Tr20^C50U^) in assay buffer [20 mM HEPES (pH 7.5), 50 mM NaCl, 50 mM KCl, 1 mM MgCl_2_,) were plunge-frozen using a Vitrobot Mark IV (Thermo Fisher Scientific) set to 100% humidity and 4 °C with the following blotting parameters: wait time 1 s, blot force 5 u and blot time 3 s. The datasets were collected on a 300 keV Titan Krios G3i (Thermo Fisher Scientific, Solaris, Poland) equipped with a Gatan BioQuantum energy filter and a Gatan K3 direct electron detector. The under-focus range was -0.9 to -2.1 µm for n-Tr20^C50U^ and -0.9 to -1.8 µm for n-Tr20. At each position, a total of 40 frames were collected that accumulated an overall dose of 40 e*^−^*Å^2^. The respective numbers of micrographs collected for each data set can be found in the Supplementary Table **1**. The pixel size used for both datasets was 0.86 Å. (Note: The tRNAs used for cryo-EM studies included a 3’-CCA in contrast to the nanopore and SAXS studies.)

### Image processing

All micrographs were subjected to motion correction using patch motion correction in Cryosparc (version 4.6.2) with default parameters with subsequent patch CTF estimation. The micrographs were then subjected to micrograph denoising using a pre-trained model and a grayscale normalization factor of 1 for further analysis. Blob picking was performed on a subset of micrographs to pick an initial set of particles. The particles were extracted and reference-free 2D classification was used to confirm the presence of RNA particles with the expected size. After validation of the selected particles, the entire data set was subjected to blob picking with the same parameters. After initial 2D classification, those classes of particles with a tRNA-like shape or equivalent size were used to create new 3D models using *ab initio* reconstruction. The particles falling into the classes containing features were pooled and subjected to another round of ab initio reconstruction (6 classes) to finally select the subset of particles with RNA-like features. Finally, classes exhibiting RNA features were checked for the presence of junk particles by 2D classification and refined by homogeneous refinement (HomoRef) [**50**]. The FSC correlation curves were calculated for each final reconstruction and the resolution was estimated using the gold standard FSC threshold of 0.143.

## Results & Discussion

We used an ultra-thin SiN*_x_* based solid-state nanopore membrane with 100 nm SiO_2_ underlayer supported by Si [**19**] to study the conformational states of n-Tr20 and n-Tr20^C50U^ using resistive-pulse sensing (**Fig. 1a**, **1b**). To optimize the experimental conditions, we tested electrolyte salt concentrations ranging from 100 mM to 2000 mM to determine the lowest salt concentration that yields an acceptable signal-to-noise ratio (SNR) and a suitable rate of tRNA translocations. High salt concentrations are known to affect tRNA conformations [**51**, **52**], while low salt concentrations significantly reduce the translocation frequency through the nanopores. The results of our optimization studies (not shown) led to our choice of 500 mM NaCl for all subsequent experiments (unless indicated otherwise; some instances required the use of 500 mM KCl).

We and others have used native polyacrylamide gel electrophoresis (nPAGE) to investigate the presence of multiple conformations [**14**, **54**, **55**], which are distinguished based on their distinct electrophoretic mobilities. To maintain uniform conditions between different experiments in this study, we performed nPAGE experiments in MgCl_2_ concentrations ranging from 0 mM to 50 mM and in either 500 mM NaCl or KCl (**Fig. 1c**). We found that n-Tr20 and n-Tr20^C50U^, folded in the absence of Mg^2+^, migrate as a single species (labeled B2). As [Mg^2+^] is increased in the refolding buffer, n-Tr20 transitions to a single slower-migrating species (B1). Because Mg^2+^ stabilizes the tertiary structure of tRNA, B1 is likely the native fold/ensemble [**14**]. However, a very small fraction of n-Tr20 was still in B2, the non-native state/ensemble, even in 50 mM MgCl_2_ and 500 mM KCl. It is possible that interactions with potassium ions (K^+^) [**56**, **57**] account for this behavior. Unlike n-Tr20, n-Tr20^C50U^ shows at least two different mobilities even in 50 mM MgCl_2_ (B1 and B2); this trend is maintained in the presence of 500 mM NaCl or 500 mM KCl (**Fig. 1c**). Thus, only a small fraction of n-Tr20^C50U^ adopts a native-like conformation/ensemble.

For nanopore experiments, selecting the optimal nanopore size and applied bias is crucial to (i) ensure a good SNR, (ii) reduce the blockage frequency during long-term measurements, and (iii) accommodate the dimensions of the tRNA molecule. Larger nanopores tend to produce lower SNR, whereas nanopores smaller than or comparable in size to the translocating (bio)molecule may cause stretching or distortion of molecular domains, leading to incorrect inferences about the (bio)molecule structure[**58**]. Similarly, high bias conditions can also induce the unfolding of the molecule during translocation [**59**]. Thus, nanopores ranging from 3.9 ± 0.4 nm to 11.9 ± 0.6 nm were tested to identify the optimal nanopore dimensions for the analysis of n-Tr20 and n-Tr20^C50U^ (**Fig. 2**).

**Figure 2:**
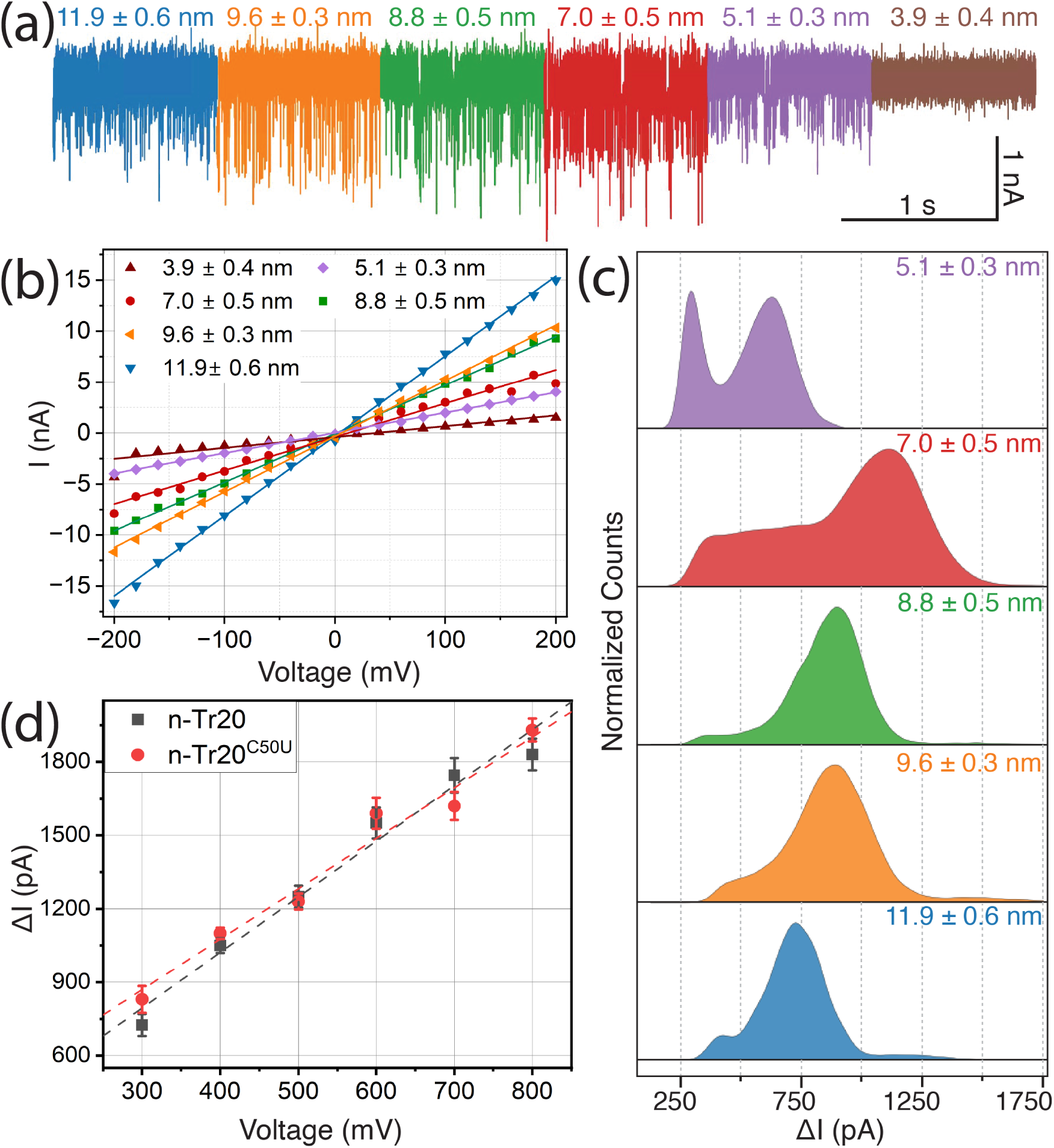
Investigation of optimal nanopore size and applied bias for tRNA experiments. (a) Representative current traces of n-Tr20 translocations through nanopores of varying diameters ranging from ∼4 to ∼12 nm. The measurements were conducted with 10 nM n-Tr20 in 5 mM Tris (pH 8) and 500 mM NaCl, with a bias of +400 mV. The corresponding open pore current-voltage traces for each pore size are shown in (b). The nanopore sizes were estimated using Equation 1 (see Methods). (c) Histograms illustrating the drop in ionic current (ΔI) when n-Tr20 molecules translocate through nanopores of varying sizes, highlighting the dependence of the current blockade results on pore diameters. (d) Observation of near-linear, voltage-dependent decrease in current (corresponding to the peak of the histogram, ΔI_peak_) for n-Tr20 and n-Tr20^C50U^. The dotted lines represent linear fits to the data.

Figure 2a shows representative ionic current traces of n-Tr20 translocations through pores of different sizes, while Fig. 2b plots the current-voltage (I-V) characteristics of these nanopores in 500 mM NaCl. The translocations were recorded at a sampling rate of 200 kHz and filtered at 35 kHz (determined as the optimal cutoff frequency [**37**]). No tRNA translocations were observed through nanopores with diameter ∼ 3.9 nm, which is not unexpected, as the simulations (using CentroidFold [**60**] and RNAComposer [**61**]) predict that the smallest dimension of the tRNA is ∼5 nm. Figure 2c displays the histograms corresponding to the drop in current (ΔI) for each nanopore size for which translocations were observed. ΔI is a measure of volumetric occlusion by the tRNA analyte during its passage through the pore. When using a 5-nm diameter nanopore, two populations were observed, centering at approximately 300 pA and 630 pA; they likely correspond to collisions of tRNA molecules with the nanopore and and back deflections (300 pA) or the smaller of two populations of tRNA (discussed more below) translocating the nanopore (630 pA). With a ∼7-nm nanopore, these populations with lower ΔI are still present. However, prevalent blockages observed suggest that many tRNA molecules have a size comparable to that of the nanopore diameter and encounter difficulty translocating through the nanopore. As the pore diameter increases from ∼7 to ∼11.9 nm, the major peak shifted to the left, a trend reflecting the increased ease of translocation by the two tRNA populations. Based on these findings, we selected nanopores of ∼8 nm for subsequent experiments. For nanopore sensing, the diameter of the pores should be close to the size of the translocating molecule for achieving optimal SNR; our choice of 8 nm which we determined empirically, is consistent with the maximum dimension of 8 nm deduced from the crystal structure of tRNA^Phe^ [**62**, **63**].

We also observed a nearly linear relationship between the applied bias and ΔI_peak_ (ΔI values at the peak of the histogram) for both n-Tr20 and n-Tr20^C50U^ (Fig. 2d). This correlation indicates that varying the applied bias does not significantly affect the experimental results. For all further experiments, we chose a bias of 400 mV as the standard operating condition.

To draw inferences on the structural ensemble of n-Tr20 and n-Tr20^C50U^, we illustrate ΔI using histograms (the corresponding scatter plots are shown in the supplementary information, **Fig. S1**). Each histogram is derived from the analysis of tens of thousands of translocation events (Fig. 3 lists the number of events). We previously showed that Mg^2+^ dictates the folds of n-Tr20 and n-Tr20^C50U^ (Fig. 1c) [**14**]. Therefore, we first investigated the effect of Mg^2+^ (included during tRNA refolding) on the nanopore translocation characteristics of n-Tr20 and n-Tr20^C50U^. Our ΔI analysis (Fig. 3a) revealed three distinguishable populations for n-Tr20 in the absence of Mg^2+^ that collapse to a single population upon the addition of 50 mM Mg^2+^(Fig. 3b) (even in 5–10 mM Mg^2+^, not shown). This observation suggests that all of the wild-type tRNA is folded to adopt a native fold/ensemble. In contrast, n-Tr20^C50U^ switches from one to three different populations when exposed to 50 mM Mg^2+^, as evidenced by the broadening of the distribution of current blockade amplitudes (Figs. 3c, 3d). Although a triple Gaussian function was used to fit the data obtained from both n-Tr20 (0 mM Mg^2+^) and n-Tr20^C50U^ (50 mM Mg^2+^), the overall distribution is narrower in the latter (see below).

**Figure 3:**
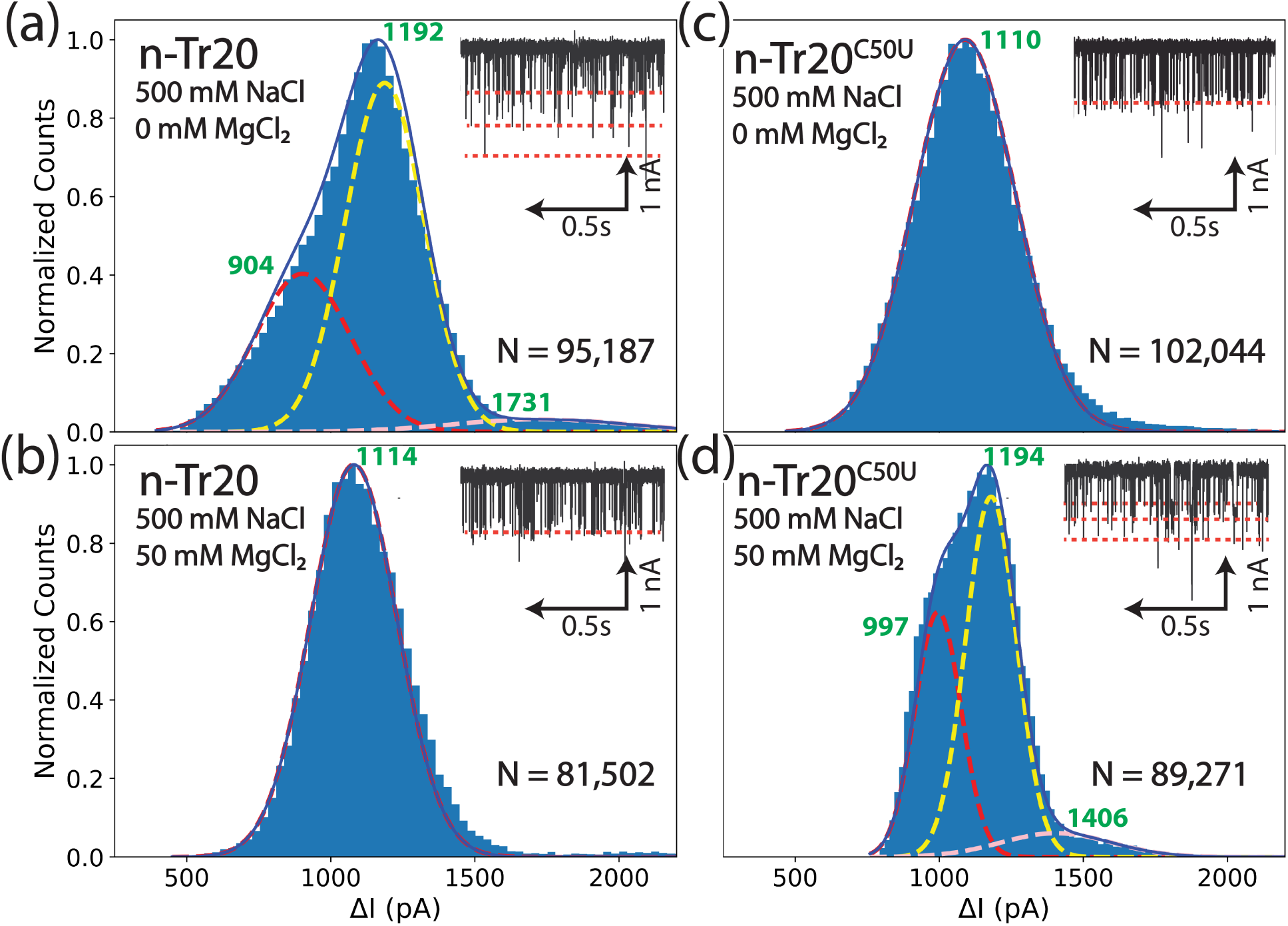
Solid-state nanopore measurements of n-Tr20 and n-Tr20^C50U^. Comparative histograms of the ionic current reduction (ΔI) observed during nanopore-sensing experiments for n-Tr20 (a, b) and n-Tr20^C50U^ (c, d). Insets display representative ionic current-time traces corresponding to translocation events for each condition indicated. N denotes the total number of events measured and used to generate the histograms. The peak positions corresponding to ΔI values are annotated at the peak maxima. n-Tr20 exhibits three distinct populations (represented by gaussian peaks) when measured in the absence of MgCl_2_, but coalesces into a single population under 50 mM MgCl_2_. In contrast, the n-Tr20^C50U^ exists as a single population in the absence of MgCl_2_, but exhibits three distinct conformations under 50 mM MgCl_2_.

Our finding of three distinct ΔI populations (n-Tr20 in 0 mM Mg^2+^), more than the two apparent states detected by nPAGE (Fig. 1c), highlights the ability of nanopore sensing to better identify the conformational ensemble in solution. The native gel mobility of each of the RNA conformers is dictated by the RNA shape/size and the size of the polyacrylamide pores, similar to the nanopore measurements that reflect the cross-sectional area of the different tRNA structural states. However, the caging effect of the native gel matrix may have narrowed the true conformational space.

We previously postulated that while n-Tr20 (in the absence of Mg^2+^) samples both native and non-native conformational ensembles, Mg^2+^ addition leads to a bias in favor of the native state. In contrast, n-Tr20^C50U^ appears to be trapped in non-native ensemble state(s), as reflected by our finding that ∼75% of pre–n-Tr20^C50U^ being intractable to 5*^′^* processing by RNase P even in 5 mM Mg^2+^ [**14**]. The ΔI histograms (Fig. 3) are consistent with this scenario. Furthermore, despite the conformational heterogeneity of n-Tr20^C50U^ in the presence of Mg^2+^, the higher ΔI observed for n-Tr20^C50U^ relative to n-Tr20 (Figs. 3c, 3d) suggests a predominantly larger-sized conformation for the former. Some of the secondary structures predicted for n-Tr20^C50U^ suggest longer dimensions compared to those for n-Tr20 (**Fig. S2**). Considering the range of potential conformations, we propose that both n-Tr20 and n-Tr20^C50U^ are capable of adopting a few different structures [**14**]. The expectation of larger conformations for n-Tr20^C50U^ aligns with cryo-EM observations (Supplementary Figures **S7, S8**).

To complement the solid-state nanopore measurements and examine the radius of gyration of n-Tr20 and n-Tr20^C50U^ under different conditions, we employed SAXS. A tRNA concentration of 0.5 mg/mL (∼20 *µ*M) was used in these studies. **Supplementary Figures, S3 and S4** presents the PRIMUS plots and the Kratky plots for n-Tr20 and n-Tr20^C50U^ in 5 mM Tris buffer (pH 8), 500 NaCl with and without 50 mM MgCl_2_. PRIMUS-based quality analysis confirmed the absence of significant aggregation or interparticle interference [**64**]. The sharply bent profile in the Kratky plots indicates that the tRNA remain folded in solution [**65**]. Using these data, we calculated the radius of gyration (*R_g_*) and found that it decreased by approximately 13% and 14% for n-Tr20 and n-Tr20^C50U^, respectively, upon the addition of Mg^2+^, reflecting structural compaction (**Supplementary Figure S5**). Furthermore, the *R_g_* values for n-Tr20^C50U^ were ∼8% higher than those for n-Tr20, with and without Mg^2+^, consistent with the predictions of our model and the nanopore measurements.

Single particle cryo-EM has emerged as a useful tool to study the dynamic three-dimensional structures of folded RNA molecules [**66**]. In addition to different larger assemblies of RNAs [**67**], we and others have recently shown that the shape of small folded RNAs, such as tRNAs, can be studied by cryo-EM [**68**–**72**]. To further substantiate the conformational differences between n-Tr20 and n-Tr20^C50U^, we prepared cryo-EM grids of the *in vitro* transcribed and refolded tRNA samples. The buffer used contained 20 mM HEPES (pH 7.5), 50 mM KCl, and 50 mM NaCl. 1 mM MgCl_2_ was added after the refolding process was complete. We collected single particle datasets (Supplementary Table **1**), analyzed detected particle stacks, and reconstructed the coulomb density maps of individual classes of molecules (Supplementary Figures **S7 and S8**). Despite the limited resolution, the results show that basically all of the ∼250,000 detected n-Tr20 particles fall into different canonical tRNA shape classes that show a high degree of structural heterogeneity. In contrast, the ∼170,000 detected particles n-Tr20^C50U^ can be classified into both L-shaped molecules and elongated particles that resemble the alternative conformation that we had postulated previously [**14**] (Supplementary Figure **S2**) and complement and confirm the distribution of conformational differences between n-Tr20 and n-Tr20^C50U^ obtained by solid-state nanopore sensing.

These low-resolution shape reconstructions from cryo-EM as well as the compaction of size measured by SAXS of n-Tr20 and n-Tr20^C50U^ are instructive. In particular, our data is consistent with the idea that mutations or modifications in RNA change the RNA structural ensemble from one distribution to another by altering fractional contributions of different conformers [**73**]. Although the cryo-EM maps obtained have too low resolution to reliably fit molecular models for the two tRNA species and it remains to be shown which specific atomic changes lead to the formation of an alternative conformation in n-Tr20^C50U^, these analyses agree with our postulates. Furthermore, the SAXS and cryo-EM data provide cross-validation and help better understand the nanopore-sensing data.

Solid-state nanopores not only detect the presence of different conformations within a sample solution but can also provide insights into the time-resolved structural changes of biomolecules under varying conditions. To explore this capability, we investigated the utility of nanopores for capturing time-resolved tRNA conformational dynamics. In these experiments, the tRNAs were re-folded in 5 mM Tris-HCl and 500 mM NaCl, and then 1, 5 or 50 mM MgCl_2_ were added just before (t=0) nanopore measurements that were taken over a 60-min period. In the presence of 1mM Mg^2+^ and 500 mM NaCl, n-Tr20 initially (*t* ≤ 14 min) displayed three peaks (peaks 1–3) in the ΔI profile (Fig. 4a), indicative of two major conformations and one minor conformation.

**Figure 4:**
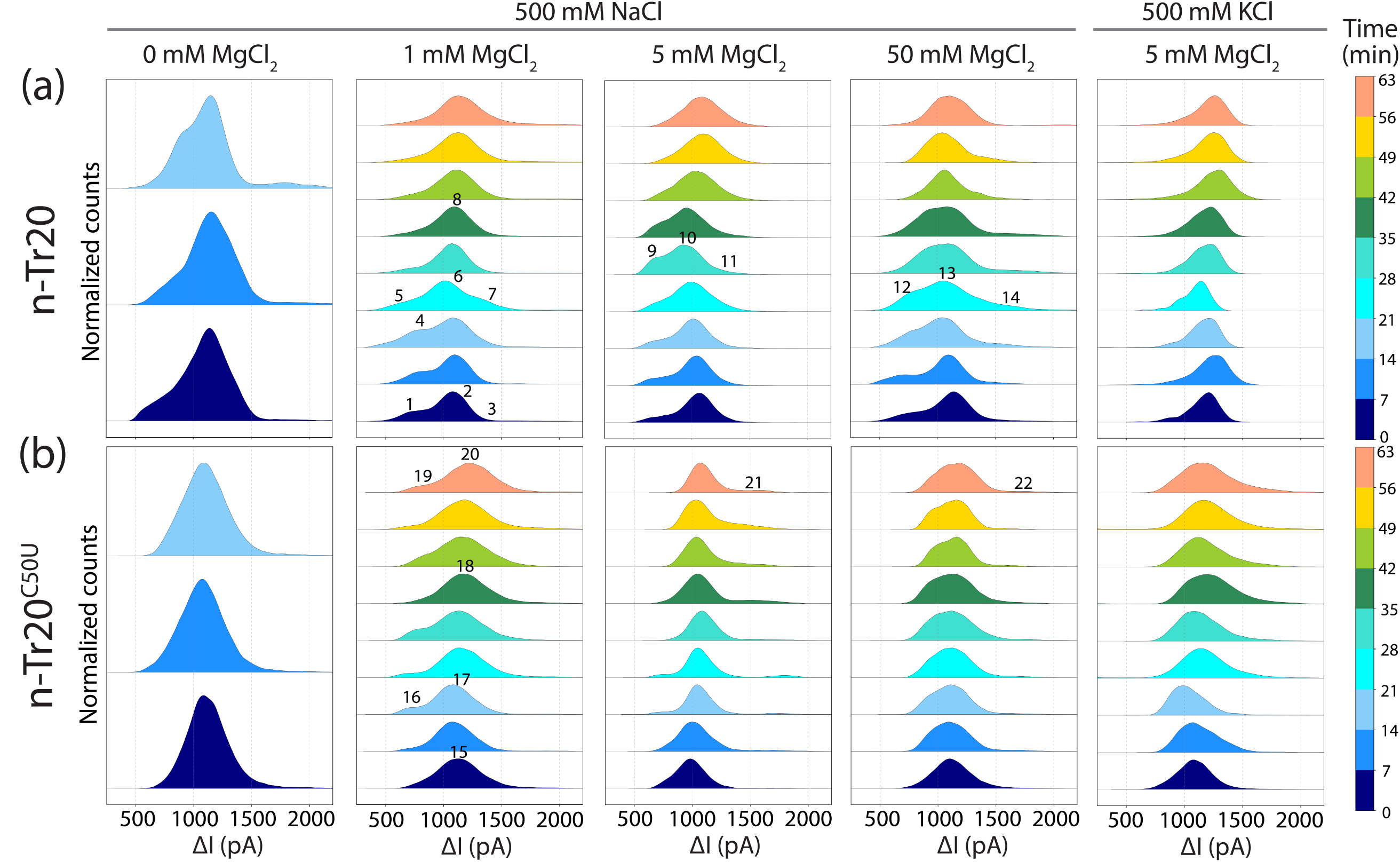
Time-resolved conformational changes of n-Tr20 and n-Tr20^C50U^. (a) Time-resolved assessments of structural alterations in n-Tr20 and n-Tr20^C50U^ in different concentrations of MgCl_2_ (as indicated) and either 500 mM NaCl or KCl as the electrolyte. Each histogram represents a 7-min measurement. Only the first three measurements (0-21 min) are shown for tRNAs without MgCl_2_, as we observed minimal changes with time. These measurements were repeated three times to confirm the reproducibility of the temporal changes.

Over time, the fraction of the peak corresponding to a lower Δ*I* value (peak 4) increased, followed by the emergence of an intermediate conformation (peak 7) and redistribution among two lower ΔI peaks (peaks 5 and 6). Gradually, these three conformations merged into a single stable conformation (peak 8). A similar pattern of metastable intermediates was observed in the presence of 5 mM and 50 mM Mg^2+^ (peaks 9–14). In agreement with our previous observation of Mg^2+^-induced structural compaction (Fig. 3), we also observe a tighter ΔI profile with increasing Mg^2+^ concentration (Fig. 4).

The conformational dynamics of n-Tr20^C50U^ (Fig. 4b) differed significantly from that of n-Tr20. At *t* ≤ 14 min, n-Tr20^C50U^ in 1 mM Mg^2+^ a single major peak was observed (peak 15). Interestingly, this single population bifurcated (peaks 16 and 17), eventually collapsing into a single population (peak 18), before redistributing into two major populations (peaks 19 and 20). Following stabilization (*t >* 40 min) in 5 and 50 mM Mg^2+^, higher-Δ*I* peaks (peaks 21 and 22) were observed, indicative of a larger-sized conformation compared to n-Tr20. These results exemplify the notion that RNA conformers can equilibrate with one another over minutes/hours [**1**]. As Mg^2+^ is increased from 1 to 5 to 50 mM, a broader range of ΔI range is apparent. These data are consistent with the view that Mg^2+^-mediated charge screening might permit local conformational relaxation and thus flexural strength [**74**].

To further investigate the effects of Mg^2+^ concentration and electrolyte chemistry (NaCl versus KCl) on tRNA conformational dynamics, we employed native-PAGE and solid-state nanopore measurements (Figs. 1c and 4). Interestingly, KCl induced structural changes more rapidly than NaCl, as evidenced by the stabilization of the ΔI values in a shorter time frame. This behavior probably reflects the ability of Mg^2+^, which is essential for tertiary folding, to outcompete loosely bound K^+^ ions more effectively [**56**, **74**]. These results demonstrate that nanopore sensing is a powerful tool for detecting conformational sampling driven by different ions.

In conclusion, our findings highlight the utility of solid-state nanopore sensing for capturing time-resolved structural changes of tRNA species. As demonstrated here, solid-state nanopore measurements, as a single-molecule method, offer near-ensemble-level characterization of RNA structures, with potential for both basic and applied research. For instance, it should be possible to assess structural homogeneity of RNA samples prepared for cryo-EM studies or as therapeutics for different diseases. Because solid-state nanopore sensing is particularly effective under high-salt conditions, it is well-suited for studies of biomolecules from halophiles or those that require salt for stabilizing functional folds. Also, even at moderate throughput, nanopore sensing-based screening approaches could help identify small molecules that alter RNA conformational equilibria. As a first step in our efforts to build a picture of the conformational ensembles sampled by a tRNA under different solution conditions, we complemented nanopore measurements with native PAGE analysis, SAXS studies and cryo-EM analysis. Looking ahead, coupling time resolved-nanopore sensing, SAXS and cryo-EM offer exciting prospects for obtaining structural information over wide-ranging time and length scales. We are buoyed by the prospects of using nanopores for illuminating the “dark space” in RNA folding, but we recognize that this potential will be realized only when different conformers identified in ΔI histograms are efficiently captured and analyzed using smFRET, native mass spectrometry, or structure-probing methods.

## Acknowledgement

Part of this work was performed on the BioSAXS beamline at the Australian Synchrotron, ANSTO. **SD** acknowledges AINSE PGRA and Australian Government RTP. **LL** and **VG** acknowledges support from the National Institutes of Health (NS-096600 to Susan Ackerman and VG), the American Heart Association (23IPA1054097 to Susan Cole and VG), and the Behrman Research Fund (to VG). **NB** and **VG** are grateful for support from a OSU PRE Accelerator Award. **PK**, **BK**, and **NB** acknowledge the ANU Grand Challenge ‘Our Health in Our Hands’ (OHIOH). Research funding for this study was provided by the European Research Council (ERC) under the European Union’s Horizon 2020 research and innovation program grant No. 101001394 (**SG**). The work is supported by the Polish Ministry and Higher Education project: “Support for research and development with the use of research infrastructure of the National Synchrotron Radiation Centre SOLARIS” under contract nr 1/SOL/2021/2. The authors gratefully acknowledge the Polish high-performance computing infrastructure PLGrid (HPC Center: ACK Cyfronet AGH) for providing computer facilities and support within the computational grant no. PLG/2023/016097 (**RM**).

## Notes

### Competing Interest Statement

The authors have declared no competing interest.

### Summary of Updates

The revised manuscript has updated figures and SI information. This manuscript has now been accepted in Nucleic Acids Research

