## Supplementary Information for "Solid-state nanopore sensing reveals conformational changes induced by a mutation in a neuron-specific tRNA^Arg^"

* Corresponding authors

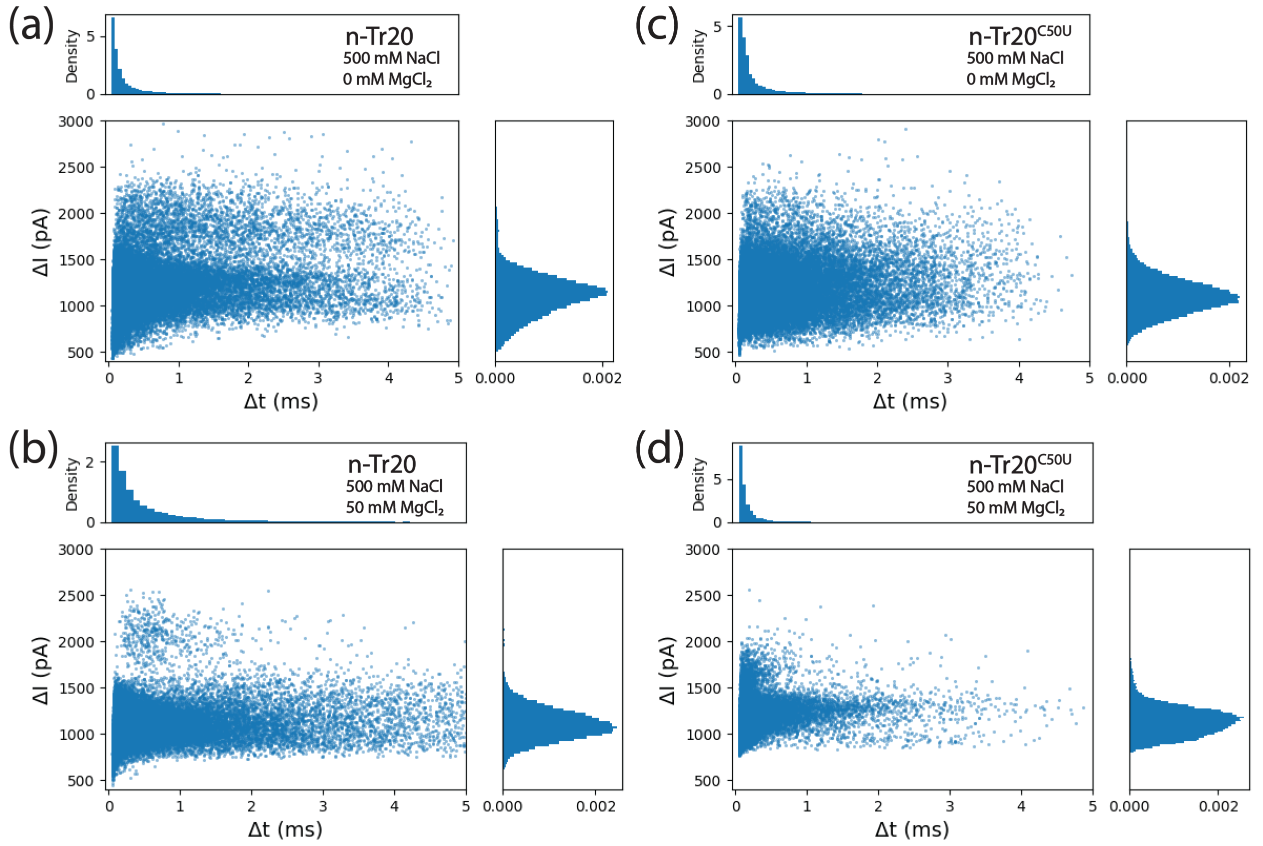


**Supplemental Figure S1.** Comparative scatter plots of the ionic current reduction (ΔI, pA) as a function of translocation time (Δt, ms). These parameters were measured during solid-state nanopore-sensing experiments with (a,b) n-Tr20 and (c,d) n-Tr20$^{C50U}$ under the conditions listed. Corresponding histograms for both ΔI and Δt are also shown on the right.

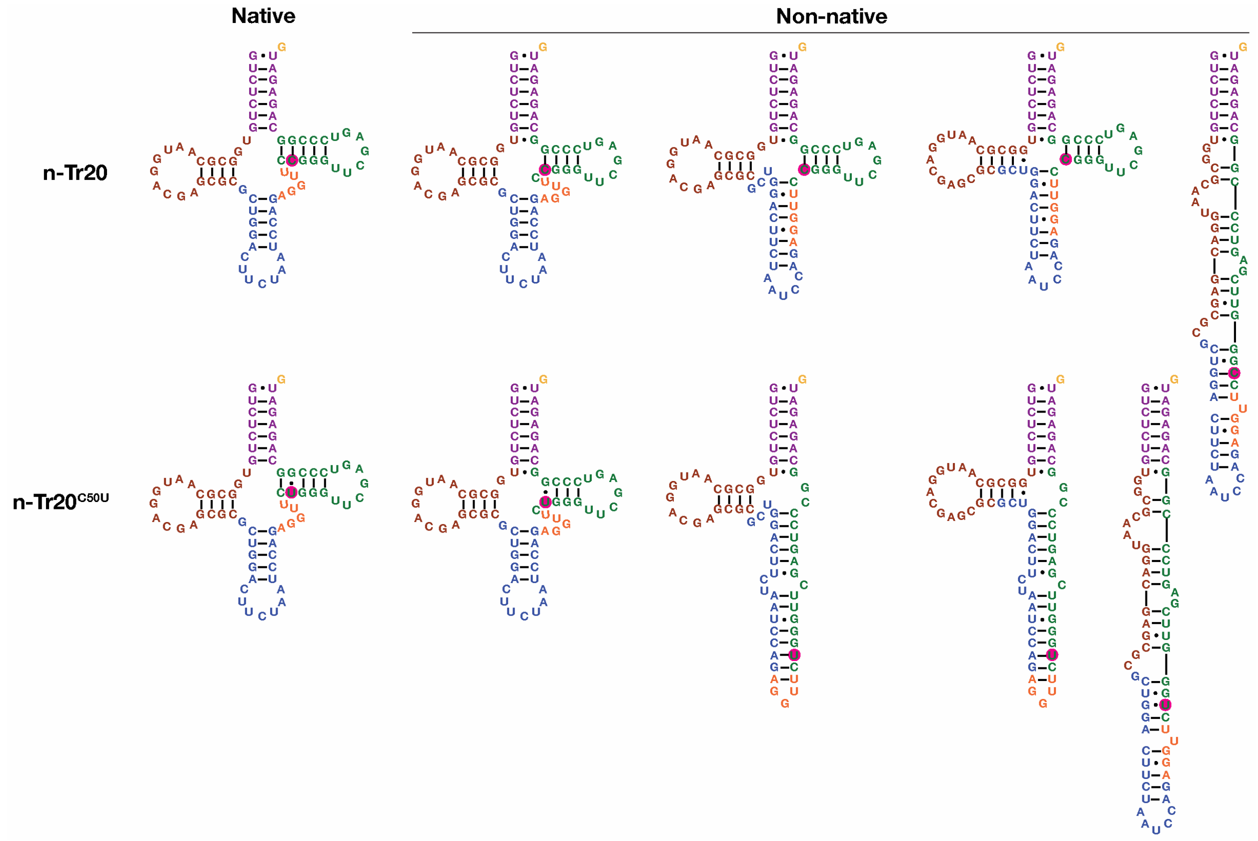


**Supplemental Figure S2.** Possible non-native conformations for n-Tr20 and n-Tr20$^{C50U}$. These non-native conformations lack the canonical "elbow" present in the native fold. These structures were predicted using the Bioinformatics Web Server for RNA (http://rtools.cbrc.jp) and manually drawn in Adobe Illustrator.

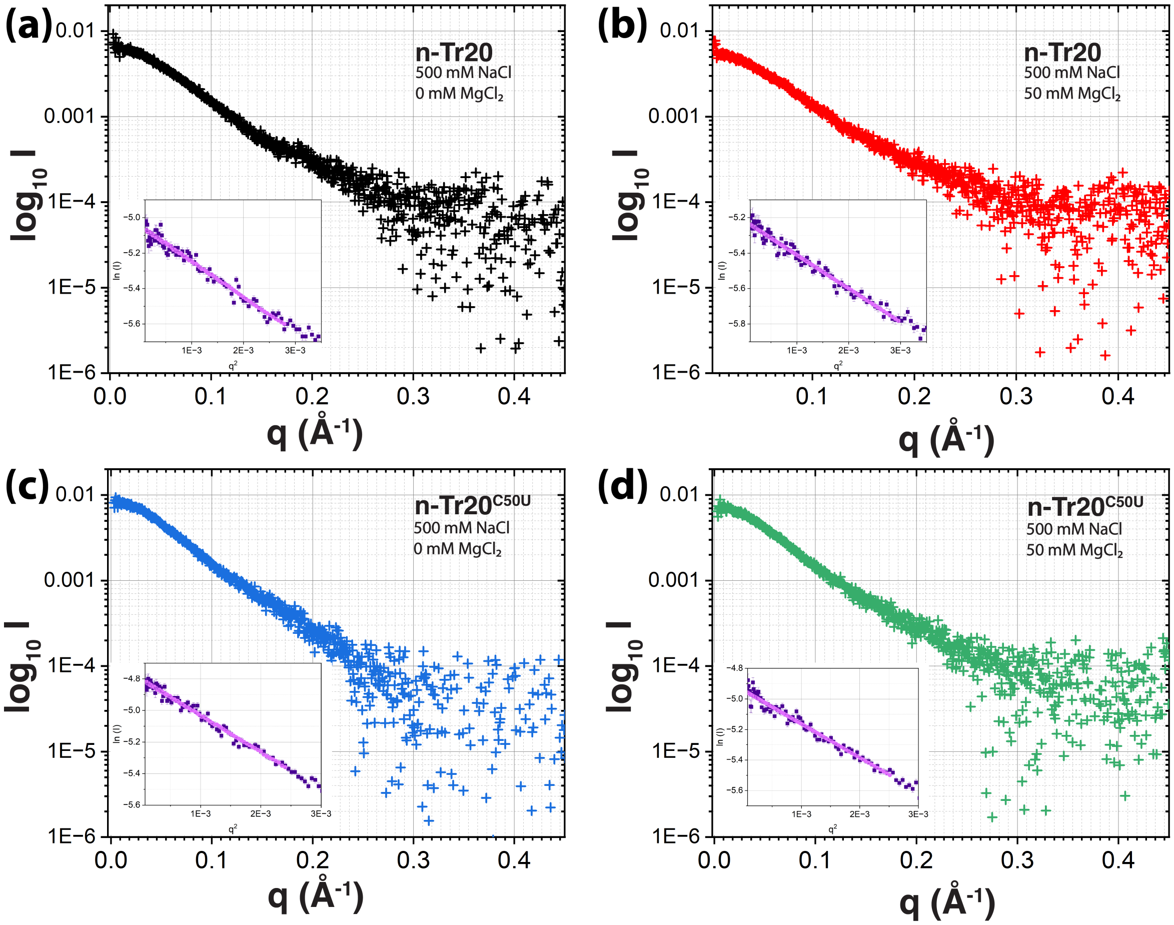


**Supplemental Figure S3.** SAXS scattering patterns plotted as $\log_{10} I$ as a function of $q$ for n-Tr20 and n-Tr20$^{\text{C50U}}$ tRNAs. (a) n-Tr20 in 500 mM NaCl, (b) n-Tr20 in 500 mM NaCl + 50 mM $MgCl_2$, (c) n-Tr20$^{\text{C50U}}$ in 500 mM NaCl, and (d) n-Tr20$^{\text{C50U}}$ in 500 mM NaCl + 50 mM $MgCl_2$. Inset: Guinier plot to determine the Radius of Gyration ($R_g$). Solid lines show the linear fits.

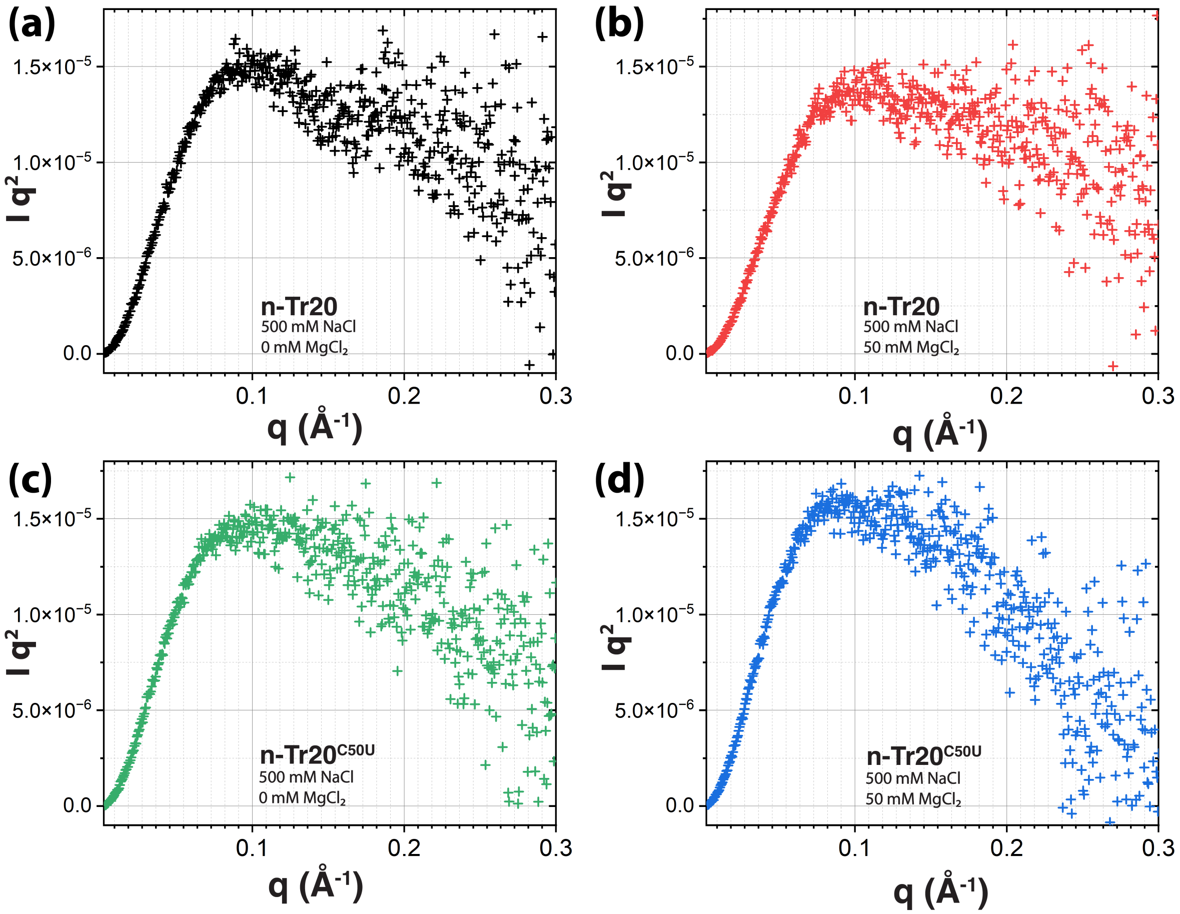


**Supplemental Figure S4.** Kratky plots ($Iq^2$ as a function of $q$) showing SAXS data for n-Tr20 and n-Tr20$^{C50U}$ RNAs under different ionic conditions. (a) n-Tr20 in 500 mM NaCl, (b) n-Tr20 in 500 mM NaCl + 50 mM $MgCl_2$, (c) n-Tr20$^{C50U}$ in 500 mM NaCl, and (d) n-Tr20$^{C50U}$ in 500 mM NaCl + 50 mM $MgCl_2$.

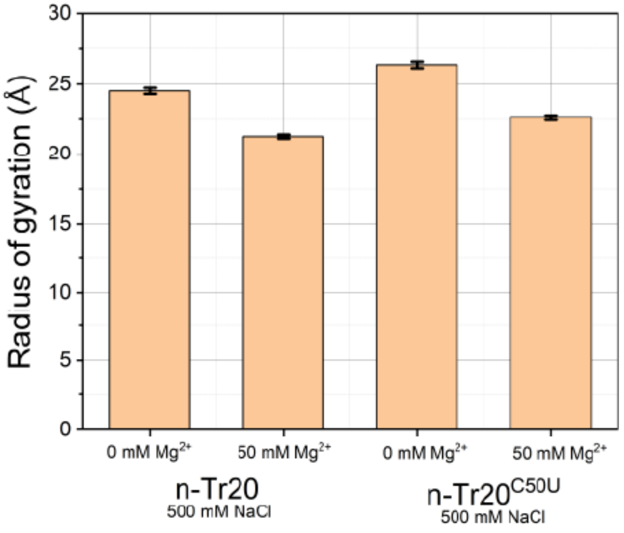


**Supplemental Figure S5.** Bar graph summarizing the radius of gyration ($R_g$) for the different samples, determined from the linear region of $\ln(I)$ vs. $q^2$ using Guinier analysis.

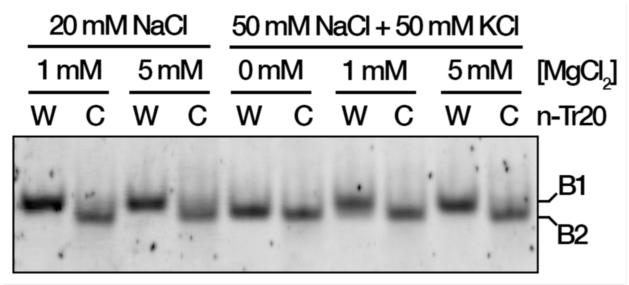


**Supplemental Figure S6.** Native PAGE analysis of n-Tr20 (W) and n-Tr20$^{C50U}$ (C) refolded in 20 mM HEPES (pH 7.5) and the indicated concentrations of $MgCl_2$ with either 20 mM NaCl (condition used in [1]) or 50 mM NaCl + 50 mM KCl (used for cryo-EM, this study). The tRNAs separated into two main bands, denoted as B1 and B2 (see text for details). It is likely that these bands represent conformational ensembles and not distinct species.

# n-Tr20

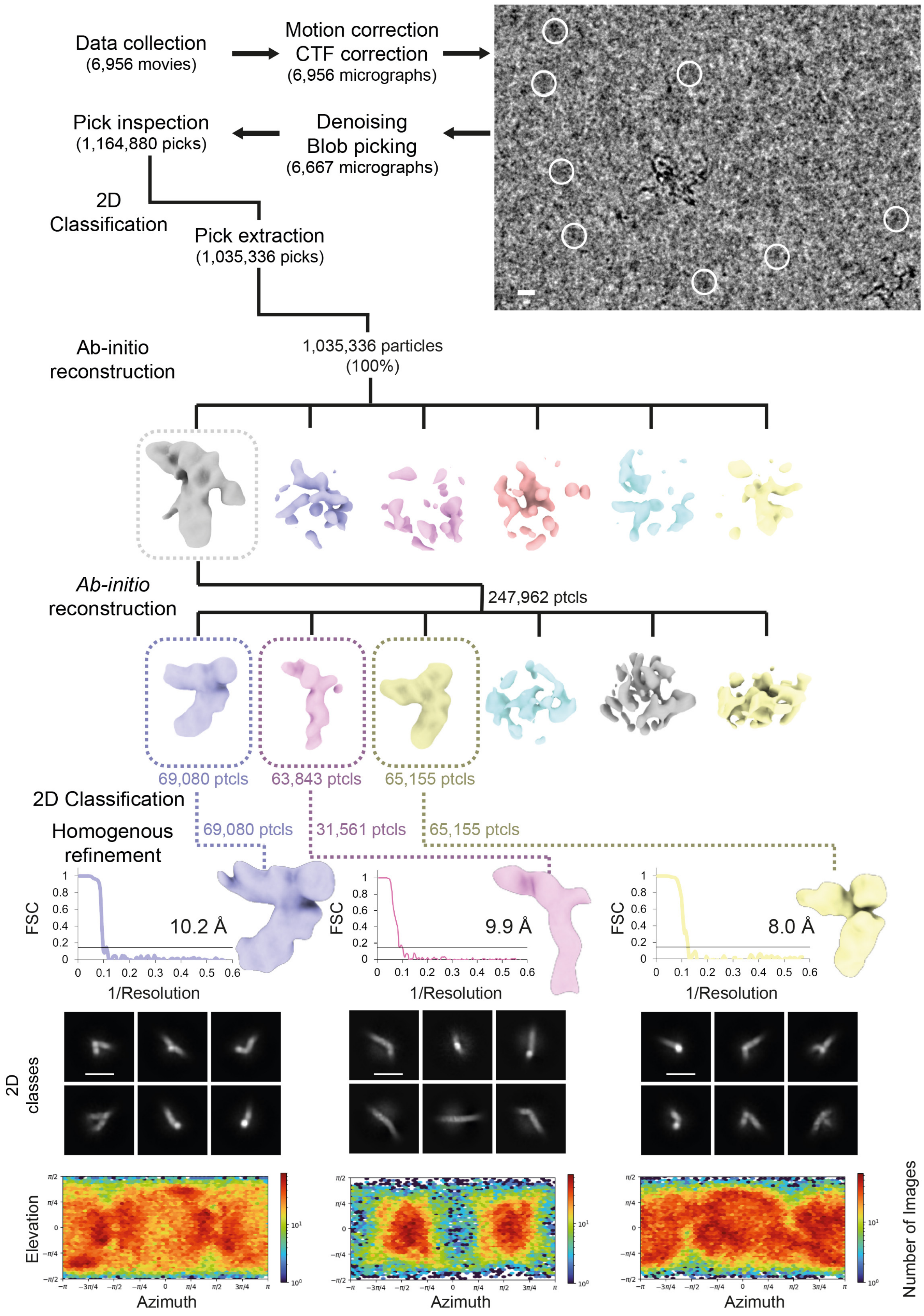

Data collection
(6,956 movies)
Motion correction
CTF correction
(6,956 micrographs)
Pick inspection
(1,164,880 picks)
Denoising
Blob picking
(6,667 micrographs)
2D
Classification
Pick extraction
(1,035,336 picks)
Ab-initio
reconstruction
1,035,336 particles
(100%)
Ab-initio
reconstruction
247,962 ptcls
69,080 ptcls
63,843 ptcls
65,155 ptcls
2D Classification
Homogenous
refinement
69,080 ptcls
31,561 ptcls
65,155 ptcls
FSC
10.2 Å
9.9 Å
8.0 Å
1/Resolution
2D
classes
Elevation
Azimuth
Number of Images

**Supplemental Figure S7. Cryo-EM reconstruction pipeline for n-Tr20.** A representative micrograph is shown (top right) together with indicated positions of the finally selected particles (white circles); scalebar = 100 Å. Representative ab-initio classes and further steps of 3D refinement are shown as well. The equally scaled cryo-EM maps of the three refined conformations with absolute numbers of particles are listed along with the Fourier Shell Correlation (FSC) plot of the final reconstruction, highlighting the nominal resolution at FSC=0.143.

## n-Tr20^C50U

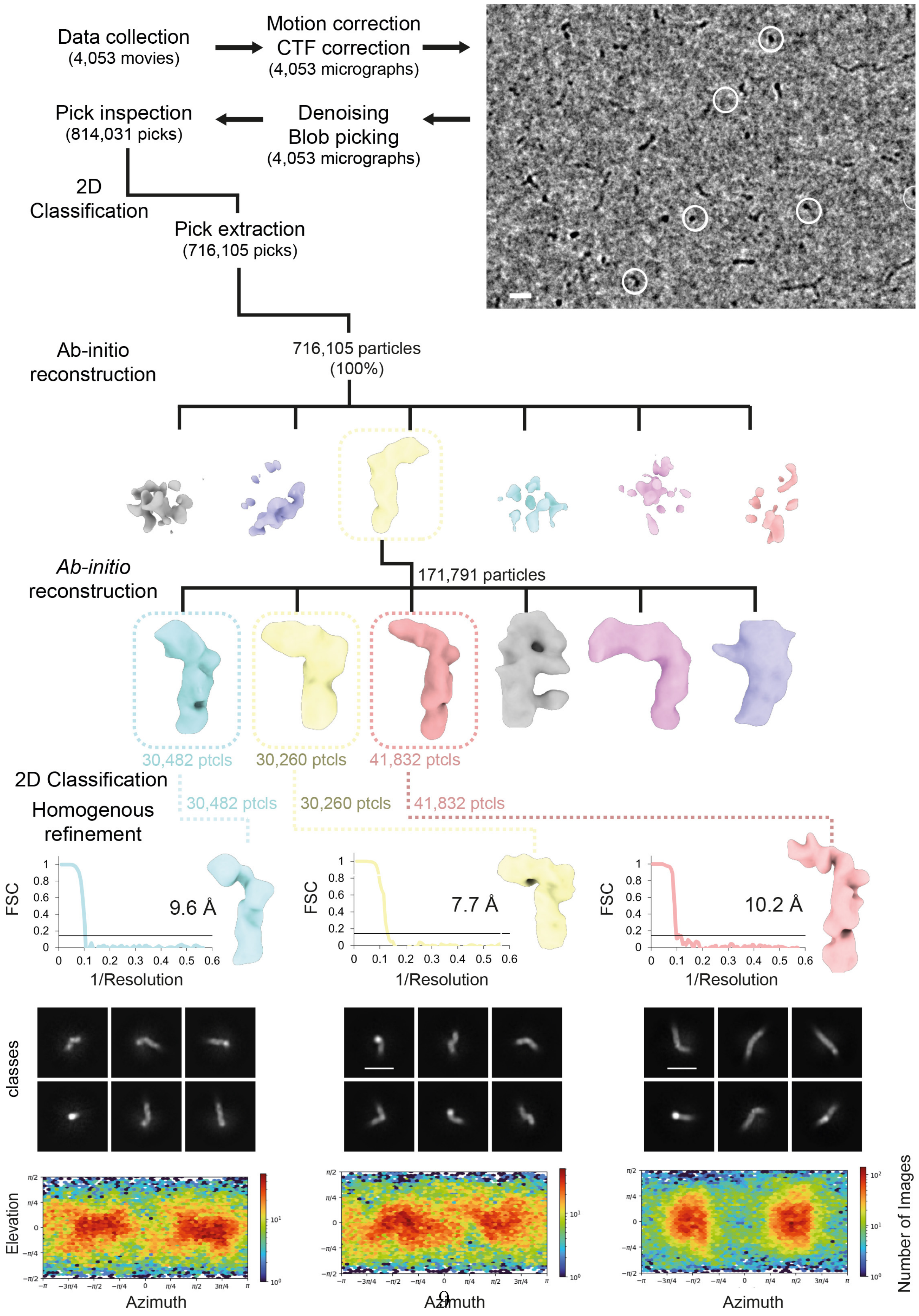
Data collection
(4,053 movies)
Motion correction
CTF correction
(4,053 micrographs)
Pick inspection
(814,031 picks)
Denoising
Blob picking
(4,053 micrographs)
2D
Classification
Pick extraction
(716,105 picks)
Ab-initio
reconstruction
716,105 particles
(100%)
Ab-initio
reconstruction
171,791 particles
30,482 ptcls
30,260 ptcls
41,832 ptcls
2D Classification
Homogenous
refinement
30,482 ptcls
30,260 ptcls
41,832 ptcls
FSC
9.6 Å
7.7 Å
10.2 Å
1/Resolution
2D
classes
Elevation
Azimuth
Number of Images

**Supplemental Figure S8. Cryo-EM reconstruction pipeline for n-Tr20$^{C50U}$.** A representative micrograph is shown (top right) together with indicated positions of the finally selected particles (white circles); scalebar = 100 Å. Representative ab-initio classes and further steps of 3D refinement are shown as well. The equally scaled cryo-EM maps of the three refined conformations with absolute numbers of particles are listed along with the Fourier Shell Correlation (FSC) plot of the final reconstruction, highlighting the nominal resolution at FSC=0.143.

**Supplementary Table S1.** Cryo-EM data collection, refinement and validation statistics.

| **Dataset** | **n-Tr20** | | | **n-Tr20$^{\mathbf{C50U}}$** | | |
|---|---|---|---|---|---|---|
| **Data collection and processing** | | | | | | |
| Magnification | 105k | | | 105k | | |
| Voltage (kV) | 300 | | | 300 | | |
| Electron exposure ($e^{-}/Å^{2}$) | 40 | | | 40 | | |
| Defocus range ($\mu$m) | 0.9-1.8 | | | 0.9-2.1 | | |
| Pixel size (Å) | 0.86 | | | 0.86 | | |
| Symmetry imposed | C1 | | | C1 | | |
| Initial particle images (no.) | 1,035,336 | | | 716,105 | | |
| EMDB | EMD-55721 | | | EMD-55720 | | |
| Final particle images (no.) | 69080 | 31561 | 65155 | 30482 | 30260 | 41832 |
| Map resolution (Å) FSC = 0.143 | 10.2 | 9.9 | 8.0 | 9.6 | 7.7 | 10.2 |
